# Varied effects of algal symbionts on transcription factor NF-κB in a sea anemone and a coral: possible roles in symbiosis and thermotolerance

**DOI:** 10.1101/640177

**Authors:** Katelyn M. Mansfield, Phillip A. Cleves, Emily Van Vlack, Nicola G. Kriefall, Brooke E. Benson, Dimitrios J. Camacho, Olivia Hemond, Monique Pedroza, Trevor Siggers, John R. Pringle, Sarah W. Davies, Thomas D. Gilmore

## Abstract

Many cnidarians, including the reef-building corals, undergo symbiotic mutualisms with photosynthetic dinoflagellate algae of the family Symbiodiniaceae. These partnerships are sensitive to temperature extremes, which cause symbiont loss and increased coral mortality. Previous studies have implicated host immunity and specifically immunity transcription factor NF-κB as having a role in the maintenance of the cnidarian-algal symbiosis. Here we have further investigated a possible role for NF-κB in establishment and loss of symbiosis in various strains of the anemone *Exaiptasia* (Aiptasia) and in the coral *Pocillopora damicornis*. Our results show that NF-κB expression is reduced in Aiptasia larvae and adults that host certain algae strains. Treatment of Aiptasia larvae with a known symbiosis-promoting cytokine, transforming growth factor β, also led to decreased NF-κB expression. We also show that aposymbiotic Aiptasia (with high NF-κB expression) have increased survival following infection with the pathogenic bacterium *Serratia marcescens* as compared to symbiotic Aiptasia (low NF-κB expression). Furthermore, a *P. damicornis* coral colony hosting *Durusdinium* spp. (formerly clade D) symbionts had higher basal NF-κB expression and decreased heat-induced bleaching as compared to two individuals hosting *Cladocopium* spp. (formerly clade C) symbionts. Lastly, genome-wide gene expression profiling and genomic promoter analysis identified putative NF-κB target genes that may be involved in thermal bleaching, symbiont maintenance, and/or immune protection in *P. damicornis*. Our results provide further support for the hypothesis that modulation of NF-κB and immunity plays a role in some, but perhaps not all, cnidarian-Symbiodiniaceae partnerships as well as in resistance to pathogens and bleaching.

## Introduction

Many corals and anemones in the phylum Cnidaria engage in mutualistic symbioses with photosynthetic intracellular dinoflagellate algae in the family Symbiodiniaceae (Muscatine and Porter, 1977; LaJeunesse et al., 2018). Different cnidarians preferentially form symbioses with distinct algal strains (Rodriguez-Lanetty et al., 2004; LaJeunesse, 2005; Coffroth et al., 2010; Hambleton et al., 2014; Wolfowicz et al., 2016), although the mechanisms determining this selectivity remain poorly understood. Disruption of coral-algal symbioses (i.e., “bleaching”), as caused by a variety of environmental factors including ocean warming, can lead to extensive coral mortality (Hoegh-Guldberg et al., 2007; Weis, 2008; Weis, 2010; Hughes et al., 2018). Thus, there has been great interest in understanding the molecular and cellular processes controlling the establishment, maintenance, and loss of algal symbiosis in cnidarians.

Corals show variable sensitivities to thermal stress based on a variety of factors including coral host genotype (Bay and Palumbi 2014; Dixon et al., 2015), symbiotic algal partner (Berkelmans and van Oppen, 2006; Oliver and Palumbi, 2011; Barshis et al., 2013), and location/thermal history (Barshis et al., 2013; Davies et al., 2018). Nevertheless, the molecular mechanisms regulating both host-symbiont selection and thermal sensitivity are not well understood. That said, the immunosuppressive host cytokine transforming growth factor β (TGFβ) has been proposed to play a role in facilitating the establishment of symbiosis in cnidarians, based on experiments with both the sea anemone Aiptasia and the coral *Fungia scutaria* (Detournay et al., 2012; Berthelier at al., 2017).

We have had an ongoing interest in immune signaling in cnidarians, specifically in the study of immune transcription factor NF-κB and its roles in pathogen infection, embryonic development, and symbiosis (Gilmore and Wolenski, 2012; Wolenski et al., 2013; Brennan et al., 2017; Mansfield et al., 2017; Williams et al., 2018). We and others have shown that the expression and activity of NF-κB are modulated during the onset of symbiosis and after loss of symbiosis in both Aiptasia (Wolfowicz et al., 2016; Mansfield et al., 2017) and the coral *Acropora palmata* (DeSalvo et al., 2010). For example, Mansfield et al. (2017) demonstrated that the establishment of algal symbiosis in naïve Aiptasia larvae is associated with a reduction in NF-κB expression and that heat-induced loss of symbiosis in adult Aiptasia is associated with increased NF-κB expression and DNA-binding activity. Furthermore, genomic analyses have shown that symbiotic cnidarians, such as the coral *Pocillopora damicornis* (Cunning et al., 2018), have extensive repertoires of immune signaling genes. Based on these data, we hypothesized that algal symbionts can modulate host NF-κB expression to effect a decrease in host immunity to allow for the establishment of symbiosis (Mansfield et al., 2017; Mansfield and Gilmore, 2019).

Herein, we have used Aiptasia and *P. damicornis* to further investigate Symbiodiniaceae-induced downregulation of NF-κB, as well as the implications of decreased NF-κB levels and symbiont type for cnidarian immunity and susceptibility to heat-induced bleaching. Our results provide further evidence for a role of NF-κB in immunity and stress responses in some, but not all, cnidarian-symbiont interactions, and further suggest that different algal types affect distinct host pathways for the establishment of stable symbioses.

## Materials and Methods

### Cnidarians and strains of Symbiodiniaceae

Adult Aiptasia used in these studies include the clonal CC7 (Sunagawa et al., 2009), H2 (Xiang et al., 2013), and PLF3 (Pringle lab stock) strains, as well as strain VW9 (a gift from V. Weis, Oregon State University) and anemones purchased from Carolina Biological. CC7 harbors clade A (genus *Symbiodinium*) algae from which the clonal, axenic strain SSA01 was derived (Bieri et al., 2016). H2 harbors a homogenous or nearly homogenous population of clade B (genus *Breviolum*) algae from which the clonal, axenic strain SSB01 was derived (Xiang et al., 2013). Other algae used include the clonal, axenic *Symbiodinium* SSA03 and clade E (genus *Effrenium*) SSE01 strains (Xiang et al., 2013; Hambleton et al., 2014), and the clade D (genus *Durusdinium*) strains Mf2.2b, A001, and Ap2 (all obtained from Mary Alice Coffroth). For routine maintenance, adult Aiptasia were kept in approximately 150 ml of 32 ppt artificial seawater (ASW) in glass or polycarbonate bowls at ~22°C with light provided by Sylvania Gro-Lux (GRO/Aq/RP) fluorescent bulbs at 20 μmol photons/m^2^/sec with a 12-h light:12-h dark cycle. Long-term aposymbiotic CC7 anemones (used in Western blots and EMSAs) had been maintained aposymbiotic for >3 years in light-proof tanks. Both symbiotic and aposymbiotic anemones were fed *Artemia* three times per week with water changes after feeding.

To generate adult Aiptasia lines harboring different algal strains, aposymbiotic CC7 anemones were infected with strain SSB01, Mf2.2b, A001, or Ap2. Infections were monitored by fluorescence microscopy for the presence and persistence of algal cells (detected by their chlorophyll fluorescence) within anemone tissue. The various symbiotic anemones were reared long-term (>1 year) under 25 μmol photons/m^2^/sec with a 12-h light:12-h dark cycle. Quantification of algal densities in these stable anemones was performed as described previously (Krediet et al., 2014) using a Guava Flow Cytometer (Millipore) to count algal cells and normalizing algal numbers to total protein measured using a Thermo Scientific Pierce BCA assay (ThermoFisher).

For experiments comparing sensitivity to bacterial infection, aposymbiotic anemones were generated by treatment of symbiotic anemones with 0.58 mM menthol as described previously (Mansfield et al., 2017). Briefly, anemones were incubated in the dark with daily changes of menthol-containing ASW for three 24-h treatment periods. After menthol treatment, anemones were placed into fresh aerated ASW and acclimated for at least two weeks before being used in assays.

To generate naturally aposymbiotic Aiptasia larvae, adult CC7 (male) and PLF3 (female) were spawned essentially as described previously (Grawunder et al., 2015). Larvae were reared in glass finger bowls in ASW until 3–4 days post-fertilization under 25 μmol photons m^−2^ s^−1^ on a 12-h light:12-h dark schedule at 27°C. Prior to infections, cultured algal cells (of strain SSA01, SSB01, SSA03, SSE01, or Mf2.2b) were rinsed three times with ASW followed by centrifugation at 3000 × g, resuspension, and counting using the Guava Flow Cytometer. Infections were performed at 50,000 algal cells/ml for 5 d in 50 ml of ASW.

The three *Pocillopora damicornis* colonies used in these studies had been maintained in common garden conditions (>5 years) at Boston University under an 8 h:16 h light/dark schedule.

### Determining NF-κB expression levels in Aiptasia larvae

For experiments assessing the ability of different algal strains to infect Aiptasia larvae and affect host NF-κB expression, larvae were infected and analyzed by fluorescence confocal microscopy, essentially as described previously (Mansfield et al., 2017). That is, larvae were fixed in 4% formaldehyde for 4 h at room temperature (RT), and after washing, samples were processed for antigen retrieval by incubating at 80°C for 30 min in 10 mM sodium citrate pH 8.5 containing 0.05% Tween. Fixed larval samples were blocked with PBS containing 0.3% Triton, 5% goat serum, and 1% BSA for 1 h at RT. Samples were then incubated overnight at RT with primary rabbit anti-Aiptasia-NF-κB antiserum (1:10,000) and mouse monoclonal anti-α-tubulin (Cell Signaling; 1:600) antiserum. Samples were next incubated with Alexa Fluor 488-conjugated goat anti-rabbit IgG secondary antiserum (Invitrogen; 1:600) and Alexa Fluor 649-conjugated goat anti-mouse IgG secondary antiserum (Invitrogen; 1:600) for 1.5 h at RT. Samples were then stained with 5 μM Hoechst 33342 (Molecular Probes). Stained larvae were imaged using channels for Alexa Flour 488 (for NF-κB), Alexa Fluor 649 (for α-tubulin), DAPI (nuclei), and mCherry (algal cell autofluorescence). Algal cells in the larvae (either in the gastric cavity or in gastrodermal cells) were counted manually by scanning through the z-planes. Corrected Total Cell Fluorescence (CTCF) of NF-κB was quantified using ImageJ. For each image, larvae were outlined using the circle tool and then area, integrated density, and mean gray value were measured; for background fluorescence, the mean gray value of an area outside the larvae was measured. CTCF was calculated by the formula (CTCF = Integrated Density – [Area × Mean gray value of background]). Statistical significance was assessed using an unpaired, twotailed t-test.

### Analysis of Aiptasia larvae treated with Transforming Growth Factor beta (TGFβ)

Approximately 300 naïve, aposymbiotic larvae were treated with 200 ng/ml of human TGFβ1 (Sigma, T7039) in 500 μl of ASW at RT for approximately 18 h. Larvae were then either fixed for immunohistochemical staining (as above) or pooled (200 larvae per treatment group) for mRNA extraction using the TRIzol reagent (Invitrogen) protocol with 10 μl of TRIzol. For qPCR, cDNA was prepared from 1 μg of RNA for each sample. RNA was combined with nuclease-free water to a final volume of 17.4 μl. cDNA synthesis was initiated by adding the following reaction components to the RNA: 5 μl of 10x RT Buffer (Applied Biosystems), 11 μl 25 mM MgCl_2_, 10 μl dNTP, 2.5 μl random hexamers, 3.1 μl MultiScribe Reverse Transcriptase (ThermoFisher), and 1 μl RNAse inhibitor (Applied Biosystems). Samples were incubated for 10 min at 25°C, 60 min at 37°C, and 5 min at 95°C. cDNA samples were used as templates for qPCR with each reaction having 2.5 μl PowerUP SYBR Green (Applied Biosystems), 0.5 μl (final concentration 200 nM) forward primer, 0.5 μl (200 nM) reverse primer, 1.25 μl dH_2_O, 0.25 μl cDNA. qPCR primers are listed in Table S1. Reactions were performed in triplicate and each primer pair included a no-template control. Reactions were carried out in standard mode on a 7900-HT Real Time PCR System (Applied Biosystems) using the following conditions: 2 min activation at 50°C, 2 min Dual-Lock DNA polymerase activation at 95°C, and 40 cycles of denaturation and annealing/extension at 95°C for 15 sec and 60°C for 1 min, respectively. Dissociation curves were generated by 15 sec incubation at 95°C, 15 sec incubation at 60°C, and then ramping from 60°C to 95°C at a 2% ramp rate.

Expression data were analyzed by the ΔΔCt method (Livak and Schmitgen, 2001). For each pooled-larvae sample performed in triplicate, average Ct values for the target gene (Aiptasia NF-κB) and reference gene (60S Ribosomal protein L10; *RPL10*) were calculated. The average Ct value for *RPL10* was then subtracted from the value for NF-κB to obtain ΔCt for control and treated groups. ΔCt values for controls were then subtracted from ΔCt for treated larvae to obtain ΔΔCt. Fold change in NF-κB mRNA was determined by 2^−ΔΔCt^. Average fold change was quantified from two experimental trials and statistical analysis was performed using an unpaired two-tailed t-test (p<0.05 for significance).

### Western blotting and electrophoretic mobility shift assays (EMSAs)

Lysates from adult Aiptasia used for EMSAs and from *P. damicornis* coral branches used for Western Blotting were prepared in AT lysis buffer supplemented with protease inhibitors (20 mM HEPES pH 7.9, 1 mM EDTA, 1 mM EGTA, 20% v/v glycerol, 1% w/v Triton X-100, 20 mM NaF, 1 mM Na_4_P_2_O_7_, 1 mM dithiothreitol, 1 mM Na_3_VO_4_, 1 mM phenylmethylsulfonyl fluoride, 1 μg/ml leupeptin, 1 μg/ml pepstatin A, 10 μg/ml aprotinin). Specifically, individual Aiptasia polyps that had been flash-frozen on dry ice/ethanol and stored at −80°C were homogenized with a plastic pestle on ice in 100 μl of AT lysis buffer, and samples were then incubated on ice for 20 min. *P. damicornis* branches were incubated in 1 ml of the buffer for 1 h at 4 °C on a rotator. In both cases, samples were next passed up and down through a 27.5-gauge needle to further disrupt tissue, and then supplemented with 4 M NaCl to a final concentration of 150 mM. Algal cells and debris were removed from samples by centrifugation at 13,000 rpm for 30 min at 4°C. Lysates containing equalized amounts of protein were then used for Western blotting or EMSA as described below.

Lysates of adult Aiptasia used for Western blotting were prepared by homogenizing whole anemones in 1.5-ml microcentrifuge tubes with a pestle in 2x SDS sample buffer (0.125 M Tris-HCl pH 8.0, 4.6% w/v SDS, 20% w/v glycerol, 10% v/v β-mercaptoethanol). For Western blotting with *P. damicornis*, lysates prepared as described above were adjusted to a final concentration of 1x SDS buffer. In both cases, samples were heated at ~95°C for 10 min followed by centrifugation at 13,000 rpm at RT for 10 min to remove insoluble material. Samples were subjected to electrophoresis in 7.5% SDS-polyacrylamide gels, and proteins were transferred to nitrocellulose overnight to ensure full transfer of high molecular weight proteins. Nitrocellulose filters were probed with our rabbit anti-Aiptasia-NF-κB antiserum (1:10,000 dilution; Mansfield et al., 2017) overnight at 4°C. (The antiserum cross-reacts with the highly similar NF-κB from *P. damicornis:* Fig. S2.) After washing, filters were incubated with antirabbit IgG horseradish-peroxidase-coupled secondary antiserum at 1:4,000 dilution for 1 h at RT, and immunoreactive bands were detected using X-ray film. To control for protein loading, filters were then stripped and probed with mouse anti-α-tubulin antiserum (Cell Signaling #3873) at 1:5,000 for 30 min at RT and then anti-mouse IgG horseradish-peroxidase-coupled secondary antiserum. NF-κB bands were normalized to α-tubulin bands, and normalized values for biological replicates were averaged. Unpaired t-tests were used to test for significant differences between control and treatment groups.

EMSAs were performed as described by Mansfield et al. (2017). Briefly, a doublestranded NF-κB-site probe (GGGAATTCCC) was end labeled with [*γ*-^32^P]-ATP (Perkin Elmer) using T4 polynucleotide kinase (New England Biolabs). For binding reactions using Aiptasia lysates in AT lysis buffer (see above), approximately 20-30 μg of total protein and 200,000 cpm of radiolabeled NF-κB-site probe were used. Reactions were carried out in binding buffer (10 mM HEPES pH 7.8, 50 mM KCl, 1 mM EDTA, 1 mM DTT, 4% w/v glycerol) for 30 min at 30°C. Samples were then electrophoresed on 5% non-denaturing polyacrylamide gels, and dried gels were subjected to autoradiography.

### Survival of Aiptasia following infection with the bacterium *Serratia marcescens*

An *S. marcescens* culture was obtained from Kim Ritchie (University of South Carolina Beaufort). Single bacterial colonies were used to inoculate 5 ml portions of Luria Broth (LB), and cultures were grown overnight at 37°C with shaking. The next day, the bacterial cells were pelleted, washed three times in ASW, resuspended in fresh ASW, and diluted to a visible light absorbance of ~0.25. Ten-ml portions of diluted bacterial cells (~1.9 x 10^8^ cells/ml) were used to infect individual anemones in single wells of a six-well tissue culture plate. Infected anemones were observed daily for 7 d, and anemones were then moved to new six-well plates containing 10 ml of fresh ASW for a two-week recovery period. Anemones were then scored as alive or dead based on tentacle regrowth and anemone movement when touched. Statistical significance of survival vs. lethality was assessed using an unpaired t-test.

### Quantification of bleaching susceptibility in Aiptasia and *P. damicornis*

For the bleaching experiments with Aiptasia, CC7 anemones–-harboring either their native symbiont or an introduced algal strain (see above)–-were heat stressed at 34°C for 6 d. Heat stress was initiated by placing tanks containing the anemones in 1 L of ASW at 27°C directly into a 34°C incubator (25 μmol photons/m^2^/sec with a 12-h light:12-h dark cycle) and allowing the water temperature to rise (over several hours) to 34°C. In each of several replicate experiments, five to eight anemones per strain were sampled at 0, 2, 4, 6, and 8 d, and algal cells were counted and normalized to total protein for each animal, as described above. To identify differences in bleaching rate among the strains, we used a generalized linear model identifying effects of anemone strain, time at 34°C, and the interaction between anemone strain and time at 34°C with these normalized algal counts. Significant pair-wise P-values from the interaction term indicate differences in bleaching rate between strains.

For *P. damicornis*, we first confirmed the species identification and characterized the Symbiodiniaceae communities. Fragments (~1 cm^3^) were collected from each of the three colonies, placed in DNA-isolation buffer (10 mM Tris-HCl pH 8.0, 100 mM NaCl, 25 mM EDTA, 0.5 % SDS, 0.1 mg/ml Proteinase K), and DNA was isolated as described previously (Davies et al., 2013). 20 ng of DNA was then used to amplify the ITS2 locus (Baumann et al. 2018) and the 18S rRNA gene (Stoeck et al., 2010). Samples were cleaned using GeneJET PCR Purification Kit (ThermoFisher), barcoded via 6 bp dual-indexing, and then pooled and sequenced at Tuft’s TUCF Genomics on an Illumina MiSeq (2 x 250 bp).

ITS2 analyses of the Symbiodiniaceae communities used the R package *dada2* (Callahan et al., 2016, with modifications outlined by Kenkel and Bay, 2018). Raw read counts ranged between 88,964 and 154,975, and after *dada2* processing, the ASV table resulted in per-sample counts ranging from 68,616 to 91,051 (File S2). Symbiodiniaceae taxonomy was assigned using the ITS2 GeoSymbio reference database (Franklin et al., 2012) using the R package *Phyloseq* (McMurdie and Holmes, 2013). The 30 most abundant ITS2 types were plotted for visualization, but all ASV counts are presented in File S2. To confirm the coral species, 18S sequences were merged and de-replicated using *usearch* (Edgar, 2010) and were assigned to taxonomic genus based on BLAST matches (Altschul et al., 1997) against nonredundant (nr) NCBI database sequences (Geer et al., 2010). All scripts are available at Github.com/NicolaKriefall/Pdam.

For the coral bleaching experiments, six 6-cm^3^ fragments were collected from each of the three colonies (18 fragments, n=9 per treatment). Fragments were randomly assigned to one of two treatments (control: 28°C; thermal stress: 32°C) such that each colony was represented in each replicate tank (n=3 per treatment). Fragments were acclimated for two days at 28°C after which thermal stress treatment (32°C) was achieved by two days of ramping at 2°C per day. Corals were maintained in 20 gallon aquaria with ASW (Instant Ocean) at 35 ppt, and light levels were standardized at 200 μmol photons m^−2^ s^−1^ using AquaIllumination Hydra TwentySix HD LED aquarium lights. Temperature and salinity were measured daily.

Two experimental trials were conducted: one for five and one for seven days, until two colonies exhibited strongly bleached phenotypes (Fig. 4A). Final photos were then obtained and fragments were immediately subsampled for RNA (200 proof ethanol) and protein (AT lysis buffer), and the remaining fragment was flash frozen in liquid nitrogen for algal density quantification. To assess bleaching, the color intensity of coral tissue was quantified from photographs of control and heated corals at the final time point of the experiment (Winters et al., 2009). Photographs contained a Coral Health Chart (CoralWatch) as a color standard. After white standardization, red, green, and blue color channel values (arbitrary units [au]) were obtained at 10 randomly placed points on each fragment. To obtain algal cell densities, ~1 cm^3^ branches were collected from each coral fragment, and tissue was extracted using an airbrush and ASW. Following tissue homogenization, algae in three 10-μl replicates were counted using a hemocytometer, and algal counts were normalized to fragment surface area using the aluminum foil method (Marsh, 1970). All statistical analyses were implemented in R (R Core Team, 2017) using the ANOVA function based on the sum of all color channels and normalized symbiont densities. Differences in colony responses across experimental treatments were evaluated for significance using post-hoc Tukey’s HSD tests.

**Figure 1.**
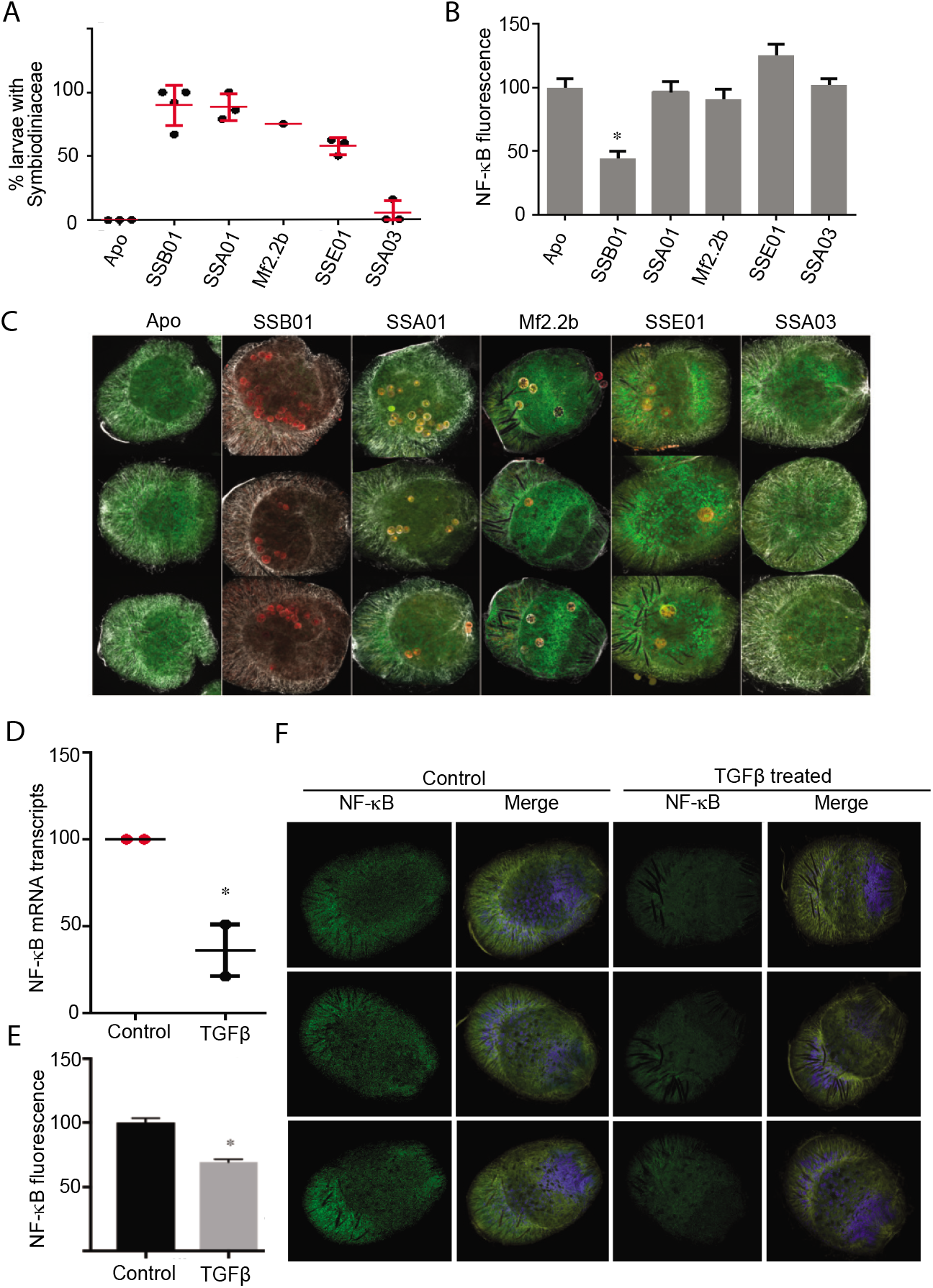
Downregulation of NF-κB expression in Aiptasia larvae by Symbiodiniaceae strain SSB01 and by treatment with exogenous TGFβ. (A) After incubation of larvae for 5 d with the indicated algal strains, the percentages of larvae containing algal cells were determined by manual counting during scans though the z-planes using confocal imaging. Each data point represents one experimental trial. (B) NF-κB protein levels were measured by quantitative immunohistochemistry of larvae exposed to the indicated algal symbionts. NF-κB fluorescence in larvae was quantified using Corrected Total Cell Fluorescence (CTCF) of confocal images as measured using ImageJ. Values represent the averages of 13 larvae for each group and are relative to the CTCF value of aposymbiotic larvae (set at 100). *= *P*<0.0001. (C) Representative immunofluorescence images of infected larvae. Green (NF-κB), red (algal cells), and white (α-tubulin) images were merged. (D-F) Aposymbiotic Aiptasia larvae were held as controls or incubated with 200 ng/ml human TGFβ ligand in ASW overnight at room temperature. (D) NF-κB and RPL10 transcript levels were measured by qPCR of cDNA from control and TGFβ-treated larvae, using RNA from 200 pooled larvae per group. Each data point is an experimental trial. Statistical significance was determined by an unpaired t-test. (E and F) Control and TGFβ-treated larvae were fixed and NF-κB fluorescence was quantified as described for (B) and (C). (E) Values are relative to the CTCF value of control uninfected larvae (set at 100). Statistical significance was determined by an unpaired t-test (n=40, *P*<0.0001). (F) Representative immunofluorescence images of control and TGFβ-treated larvae. Green (NF-κB) yellow (α-tubulin), and blue (DAPI) were merged.

### Gene expression profiling and differential expression analyses

For gene expression analyses, total RNA from corals under control conditions from the first experimental bleaching trial (n=9) was isolated and treated with DNase using an RNAqueous kit protocol (Ambion, Life Technologies). Approximately 1 μg of RNA per sample was then prepared for tag-based RNA-seq as described by Meyer et al. (2011), with several modifications to account for the transition to the Illumina sequencing platform (Dixon et al., 2015; Lohman et al., 2016). Eight of nine libraries were successfully prepared and sequenced on the Illumina HiSeq 2500 at Tuft’s TUCF Genomics yielding single-end (SE) 50 bp reads.

62.1 million raw reads were generated, with individual library counts of 5.7 to 10 million reads (mean = 7.8 million reads). Fastx toolkit was used to remove 5’-Illumina leader sequences and poly(A)^+^ tails, and sequences <20 bp in length and with <90% of bases having quality cutoff scores <20 were also trimmed. In addition, because degenerate bases were incorporated during cDNA synthesis, PCR duplicates were identified and removed from all libraries as described by Dixon et al. (2015). After quality filtering, the resulting quality filtered reads were mapped to the *P. damicornis* genome (Cunning et al., 2018) using Bowtie2.2.0 (Langmead and Salzberg, 2012), which yielded a counts file (File S1). Sample counts ranged from 1.0 to 2.1 million (mean = 1.5 million counts).

Differential gene expression analyses were performed with *DESeq2* v. 1.6.3 (Love et al., 2014) using the model: design = ~ colony. Counts were normalized, and then independent pairwise contrasts were computed across all three colony comparisons (1 vs 2, 1 vs 3, 2 vs 3). Genes identified as differentially expressed were corrected for false positives using the Benjamini and Hochberg (1995) false discovery rate (FDR) correction (adjusted *P*-value < 0.05) and genes that were consistently up- or down-regulated in each coral colony across both pairwise comparisons were determined (Files S3-8). Gene expression results for genes consistently overexpressed in colony 3 were then visualized using the R package *pheatmap*. Expression data were then rlog-normalized to broadly characterize differences in gene expression across colonies using a principal component analysis and a multivariate analysis of variance function (Adonis) using the package vegan (Oksanen et al., 2016).

### Analysis of genes upregulated in *P. damicornis* heat-resistant colony 3

Genes significantly upregulated (FDR p<0.05) in colony 3 were analyzed for upregulation in heat and other stress conditions by a literature search (see Table S2). To identify potential NF-κB binding sites in *P. damicornis* genomic sequences, the 500 bp upstream of the transcription start sites in genes of interest in the *P. damicornis* genome (Cunning et al., 2018) were examined for the presence of predicted NF-κB binding sites using Aiptasia NF-ĸB’s DNA-binding site preference (based on protein binding microarray (PBM) analyses in Mansfield et al., (2017) (sequences in File S9).

## Results

### Downregulation of NF-κB expression in Aiptasia larvae by Symbiodiniaceae strain SSB01 but not by several other strains

In a survey of wild anemones, Aiptasia was found to be symbiotic with Symbiodiniaceae strains in several genera (Thornhill et al., 2013; LaJeunesse et al., 2018). The clonal CC7 anemone line contains largely or entirely algae of strain SSA01 (*Symbiodinium* sp.) (Sunagawa et al., 2009; Bieri et al., 2016), but can also be readily populated to high levels by strain SSB01 (*Breviolum minutum*) (Xiang et al., 2013; Hambleton et al., 2014; Baumgarten et al., 2015). It cannot be stably populated with either the *Symbiodinium* strain SSA03 or the *Effrenium voratum* (clade E) strain SSE01 (Hambleton et al., 2014; Wolfowicz et al., 2016). The clonal anemone line PLF3 appears to contain a mixture of algal strains SSA01 and SSB01 (our unpublished results). To determine whether algal strains with differing abilities to establish stable symbiosis also have different effects on host NF-κB expression, we incubated Aiptasia larvae (from a cross of strains CC7 and PLF3) for 5 d with strain SSA01, SSB01, SSA03, SSE01, or the previously unexamined clade D (*Durusdinium* sp.) strain Mf2.2b. Larvae were then fixed, stained, and imaged for NF-κB expression levels.

As an initial comparison, we also counted the number of algal cells within infected larvae, based on their autofluorescence. As expected, strains SSA01 and SSB01 were found in ~90% of the larvae after a 5 d incubation, while strains SSE01 and SSA03 were found less frequently (~57% and ~5%, respectively) (Fig. 1A). Although wild Aiptasia have not been found to harbor *Durusdinium* algae, strain Mf2.2b was found in ~75% of the larvae (Fig. 1A). No algae were detected in larvae prior to exposure to the cultured algae.

As compared to aposymbiotic larvae, NF-κB expression was reduced by 52% in larvae hosting SSB01 (n=16, P<0.0001) (Figs. 1B, C). In contrast, larvae hosting native SSA01 or nonnative SSE01 or Mf2.2 showed no significant difference in NF-κB staining as compared to aposymbiotic controls. NF-κB expression was also not significantly reduced in larvae incubated with SSA03 algae, which were only rarely (<5%) observed in the larvae (Figs. 1B, C).

### Downregulation of NF-κB expression in Aiptasia larvae by exogenous TGFβ

Previous data have suggested that the cytokine TGFβ can suppress immunity in adult Aiptasia (Detournay et al., 2012), and therefore, we were interested in whether TGFβ could also affect NF-κB in Aiptasia. Indeed, treatment of aposymbiotic Aiptasia larvae with human TGFβ1 reduced NF-κB mRNA by 64% as compared to control larvae (p<0.05; Fig. 1D). Similarly, human TGFβ1 treatment of Aiptasia larvae reduced NF-κB protein staining by 31% (n=40, P<0.0001; Figs. 1E, F).

### Downregulation of NF-κB expression in Aiptasia adults by Symbiodiniaceae strains SSB01 and SSA01 but not by several *Durusdinium* (clade D) strains

To determine whether NF-κB expression is also affected in adult Aiptasia in an algal strain-dependent manner, we compared NF-κB expression levels in CC7 anemones containing their endogenous algae (largely or entirely strain SSA01) to anemones that had been fully bleached (aposymbiotic) or had been bleached and then infected with strain SSB01 or with one of three *Durusdinium* strains (Mf2.2b, A001, and Ap2).. These symbionts were chosen because adult Aiptasia infected with SSB01 have reduced NF-κB levels (Mansfield et al., 2017); strains SSA01 and SSB01 can be hosted at high levels in Aiptasia (Hambleton et al., 2014; Grawunder et al., 2015; Wolfowicz et al., 2016; Fig. 2B); and non-native strains Mf2.2b, A001, and Ap2 can establish stable symbioses, albeit at reduced population densities (Figs. 1B, C). By Western blotting, we found that CC7-SSB01 and CC7-SSA01 had reduced levels of NF-κB as compared to aposymbiotic Aiptasia (Fig. 2A). In contrast, CC7 infected with Mf2.2b, A001, or Ap2 had NF-κB levels that were similar to those in aposymbiotic anemones (Fig. 2A).

**Figure 2.**
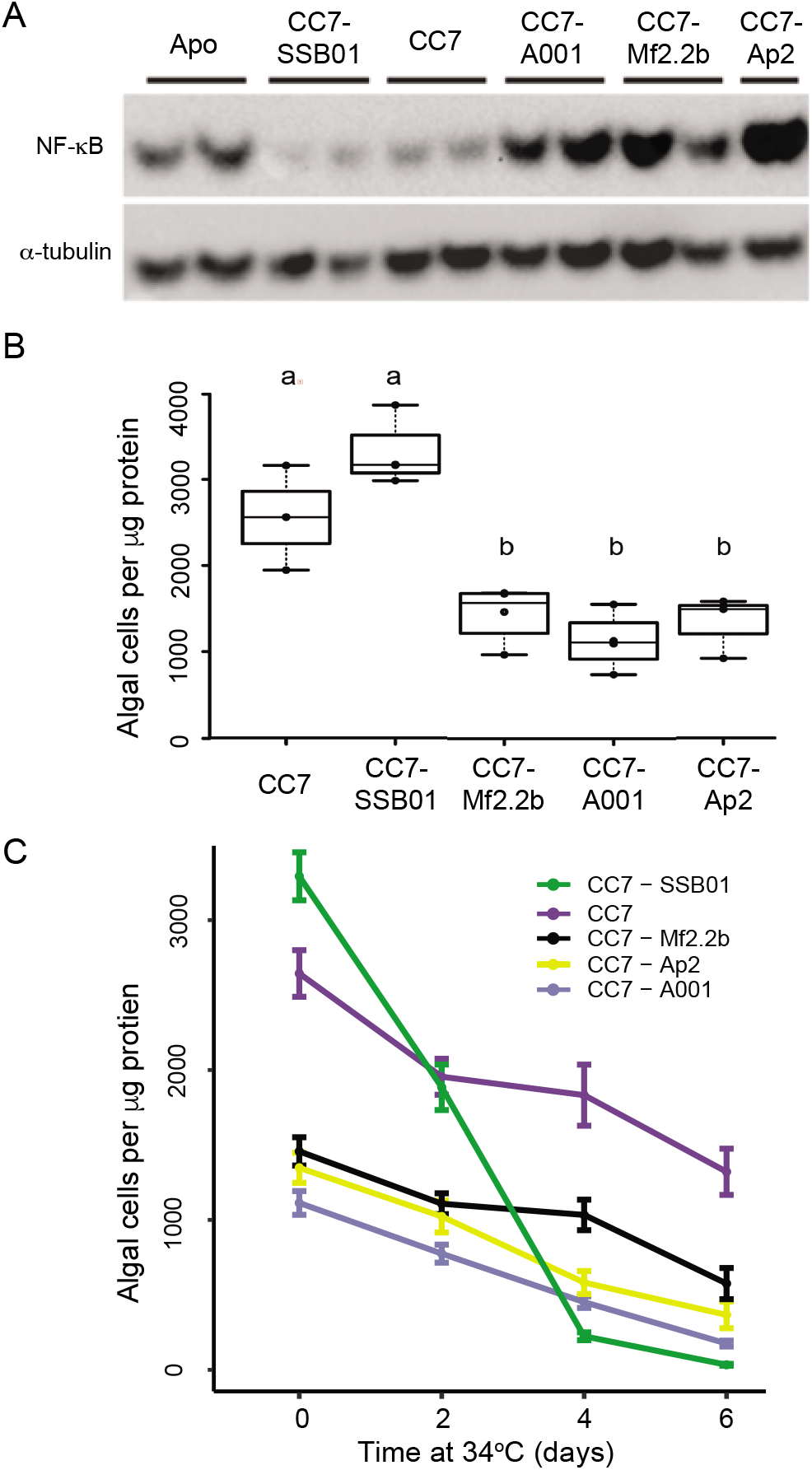
Downregulation of NF-κB expression in Aiptasia adults by Symbiodiniaceae strains SSB01 and SSA01, but not by three *Durusdinium* strains. (A) Western blotting using anti-NF-κB and anti-α-tubulin antisera of whole-animal extracts from aposymbiotic CC7 Aiptasia (Apo), CC7 animals hosting their endogenous algal symbionts (CC7), or CC7 that had been rendered aposymbiotic and then populated to steady-state levels with the *Breviolum* strain SSB01 or one of the *Durusdinium* strains (A001, Mf2.2b, and Ap2). (B) Algal population densities were determined in the same anemone strains as in (A). Box-and-whiskers plots are shown for three (CC7, CC7-Ap2) or four (CC7-SSB01, CC7-Mf2.2b, CC7-A001) replicate experiments with five to eight animals per experiment. ANOVA Tukey post-hoc tests were used to evaluate the differences between populations; those marked by the same letter were not significantly different. For each comparison of CC7 to one of the *Durusdinium-containing* strains, *P*<0.05; for each comparison of CC7-SSB01 to one of the *Durusdinium-containing* strains, *P*<0.0001. (C) Rates of bleaching under thermal stress at 34°C were evaluated for the same anemone populations as in (A) and (B). Means +/-SEMs are shown for replicates as in (B). (Note that the data for time 0 are the same as those shown in B.) ENDO, CC7 animals hosting their endogenous algal symbionts.

To determine whether thermal tolerance correlates with pre-stress NF-κB or symbiont levels across our Aiptasia strains, we subjected these five symbiotic Aiptasia strains to heat stress at 34°C (Fig. 2C). We found that CC7-SSB01, with low levels of NF-κB and high levels of algae, bleached faster than all other strains tested (each pair-wise GLM interaction P < 1 x 10^−16^). In contrast, CC7 with its endogenous algae (SSA01) or harboring any of the *Durusdinium* strains they bleached at similar rates (Fig. 2C), even though they differed markedly in both steady-state algal population levels and NF-κB levels (Figs. 2A, B). Thus, neither steady-state algal population densities nor NF-κB levels appeared to correlate with susceptibility to thermal bleaching in Aiptasia.

### Increased resistance to a bacterial pathogen of aposymbiotic (high NF-κB) relative to symbiotic (low NF-κB) Aiptasia

Due to the broad role of NF-κB in immunity in higher metazoans (Gilmore et al., 2016), we hypothesized that NF-κB activity would be positively correlated with immune competence in Aiptasia. To test this, we first compared NF-κB DNA-binding activity by electrophoretic mobility shift assay (EMSA) in lysates of adult CC7 Aiptasia that were either aposymbiotic or symbiotic with algal strain SSB01. Consistent with our measurement of NF-κB protein levels (Fig. 2A), the aposymbiotic anemones had approximately three-fold more NF-κB DNA-binding activity than symbiotic anemones (Figs. 3A, B). This result is similar to the reduced NF-κB DNA-binding activity that we previously found with H2-SSBO1 anemones (Mansfield et al., 2017).

**Figure 3.**
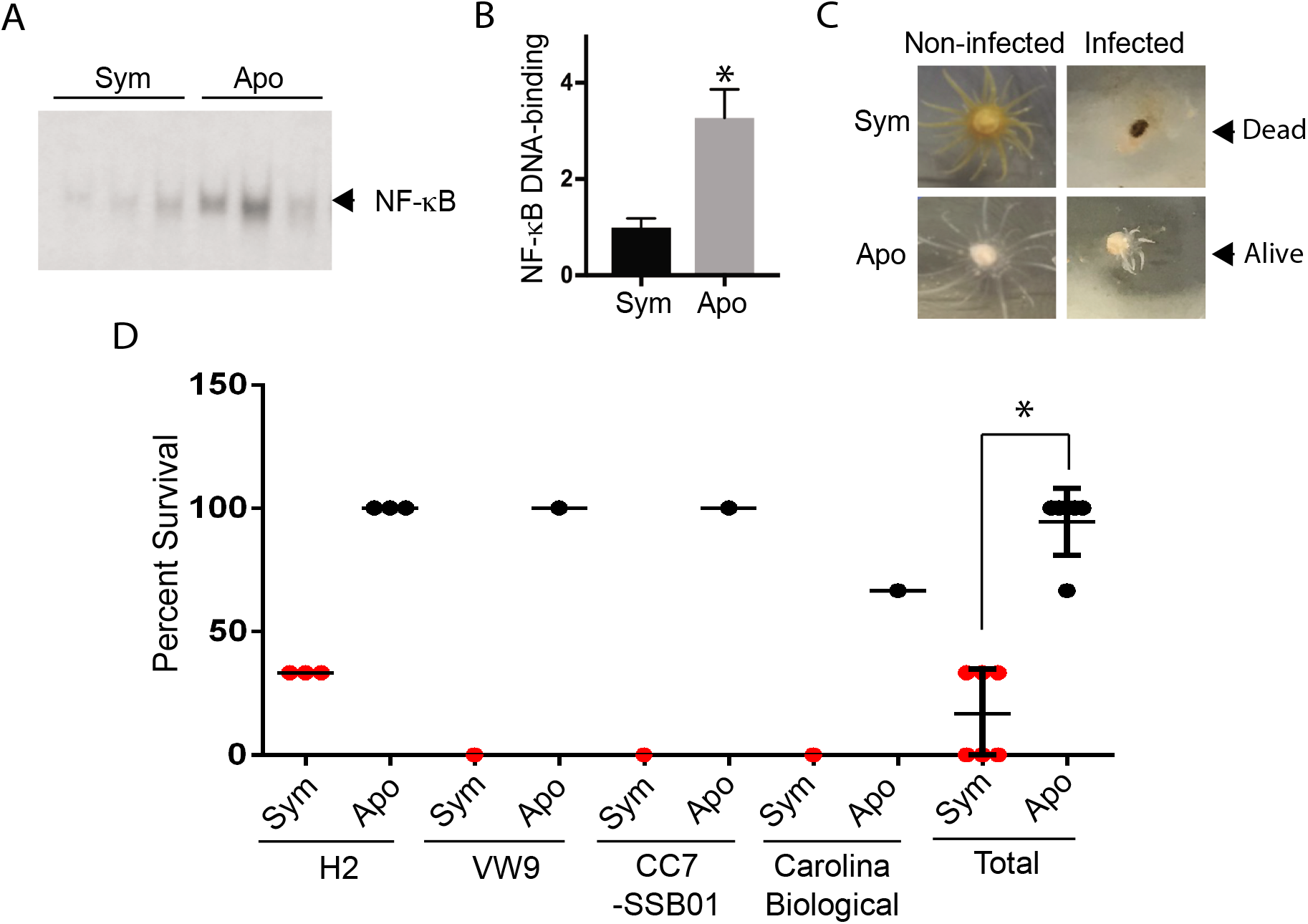
Increased resistance to a bacterial pathogen of aposymbiotic (high NF-κB) relative to symbiotic (low NF-κB) Aiptasia. (A) NF-κB DNA-binding activity was assessed by EMSA using lysates of three adult symbiotic CC7-SSB01 (Sym) or aposymbiotic CC7 (Apo) anemones. DNA-binding reactions were performed with equal amounts of protein for all biological replicates. The NF-κB DNA-binding complex is indicated by the arrow. (B) Bands corresponding to the NF-κB DNA-binding complexes in (A) were quantified using ImageJ. Values were averaged per group and then normalized to the mean value (set at 1.0) for the three bands detected with symbiotic animal extracts. (C) CC7-SSB01 (Sym) and aposymbiotic (Apo) Aiptasia were incubated with *Serratia marcescens* for one week and were then allowed to recover in ASW for two weeks at which time viability was assessed (see Materials and Methods). Representative images are shown of control, non-infected anemones (left) and infected anemones after the two-week recovery period (right); an anemone that did not survive infection (top, right), and an anemone that did survive (bottom, right) are shown. (D) Percent survival of *S. marcescens*-infected anemones for the indicated symbiotic/aposymbiotic pairs. Each data point is an experimental trial (as in C) with three anemones. The statistical significance of the cumulative Sym versus Apo data (n=18 for each group) was evaluated using an unpaired t-test (*P*<0.0001).

We next compared the sensitivities to bacterial infection of four strains of symbiotic Aiptasia relative to their aposymbiotic counterparts (generated by menthol treatment; see Materials and Methods). All anemones were challenged with. *S. marcescens*, a bacterial pathogen that causes white pox disease in wild coral (Sutherland et al., 2011) and lethality in Aiptasia (Krediet et al., 2014). Anemone survival was assessed by observing the presence of extended tentacles (Fig. 3C) and movement in response to touch by a plastic pipette. For each anemone strain, the aposymbiotic animals showed better survival than the symbiotic animals (Fig. 3D). When aggregated over all anemone strains, the difference in survival of anemones was statistically significant: ~95% of aposymbiotic Aiptasia survived infection with *S. marcescens*, whereas only ~17% of symbiotic Aiptasia survived (n=18, P<0.0001; Fig. 3D). These results demonstrate a positive correlation between increased NF-κB levels and an increased ability to survive pathogen challenge across several strains of Aiptasia.

### Higher NF-κB levels and greater thermotolerance in a Pocillopora damicornis colony hosting *Durusdinium* as compared to colonies hosting *Cladocopium* symbionts

We next investigated whether a thermosensitive coral also exhibits differences in NF-κB levels when hosting different genera of Symbiodiniaceae. For these experiments, three independent colonies of *P. damicornis* that had been maintained in common garden laboratory conditions for >5 years, were used (Fig. 4A). 18S amplicon sequencing confirmed that all three colonies belonged to the same *Pocillopora* species, as the top BLAST hit for all three colonies was “*Pocillopora damicornis* partial 18S rRNA gene”, with a 100% query cover, sourced from Arrigoni et al. (2017) (accession number: LT631138). Nevertheless, these colonies exhibited distinct gene expression profiles as demonstrated by principal component analysis of all transcripts (Fig. 5), suggesting that these colonies are genetically distinct. Algal cell counts showed that the levels of symbionts in the three colonies were not significantly different (Fig. 4B). Metabarcoding of the ITS2 locus showed that colonies 1 and 2 were dominated by *Cladocopium* (previously clade C) species, whereas colony 3 hosted primarily *Durusdinium* species (Fig. 4C).

**Figure 4.**
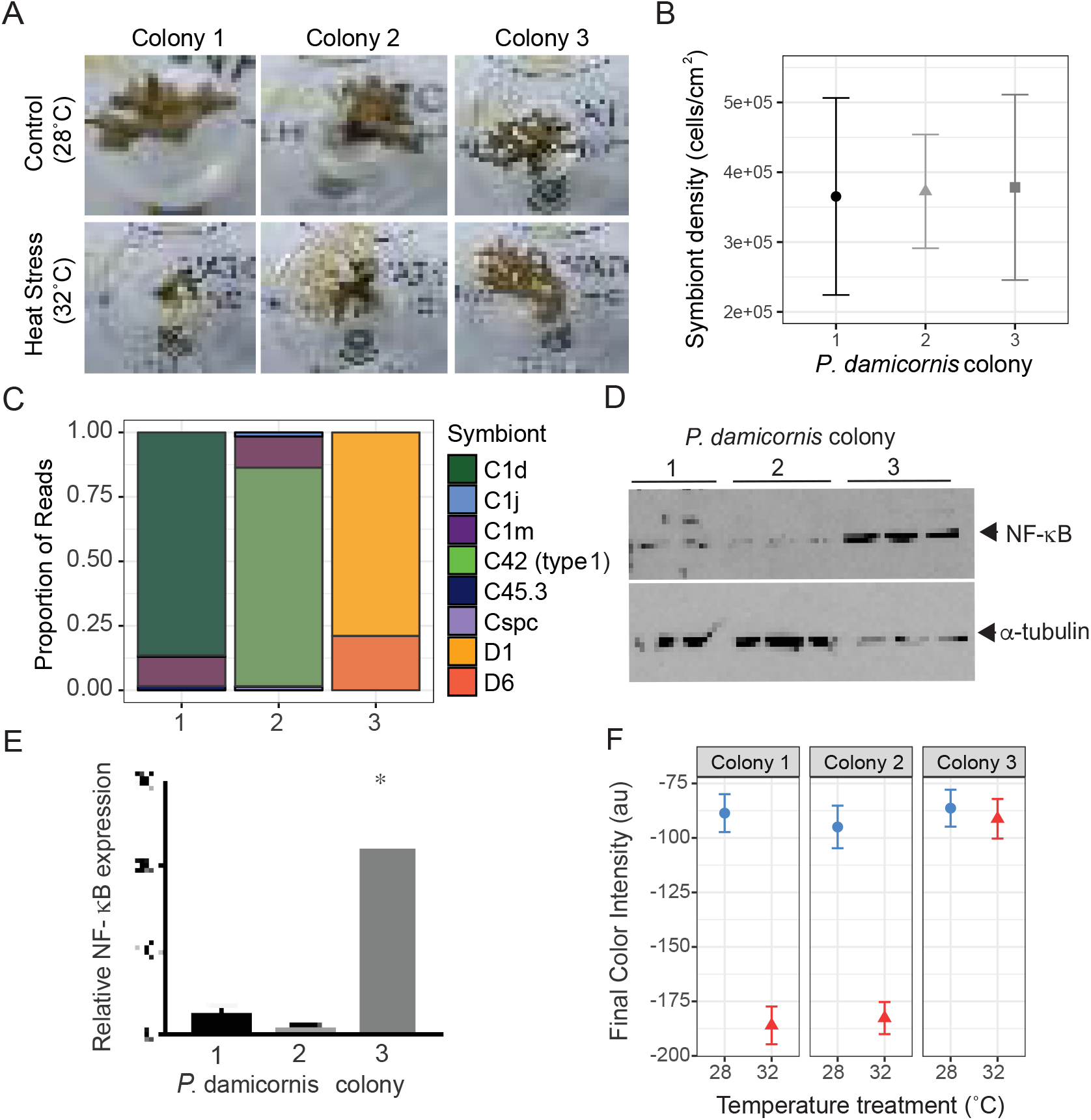
Higher NF-κB levels and greater thermotolerance in a *Pocillopora damicornis* colony hosting *Durusdinium* than in colonies hosting *Cladocopium* symbionts. Three colonies of *P. damicornis* that had been kept in laboratory-based common-garden conditions were characterized for their algal symbionts, NF-κB levels, and sensitivities to bleaching under heat Total algal population densities in the colonies at 28°C were determined as described in Materials and Methods. (C) Algal symbiont communities were characterized by metabarcoding of ITS2, and the relative abundance of each ITS2 type was plotted as the proportion of reads per sample. (D) Samples from the colonies held at 28°C for 5 days were lysed, and samples were subjected to Western blotting for NF-κB (top) and α-tubulin (bottom).). (E) NF-κB bands (as in D) were quantified in ImageJ, and intensities were normalized to α-tubulin band intensity values. Values are averages of three biological replicates from each of two experimental trials (thus n=6, except for colony 1, which had one NF-κB band below the detection limit) and are expressed relative to that for colony 2 (set at 1.0). Statistical significance was evaluated using an unpaired t-test (colony 1 vs. colony 3, *P*=0.005). (F) Tests of sensitivity to thermal bleaching. In each of two trials, six fragments from each colony were randomly assigned to one of two treatments, either continued incubation at 28°C (control) or incubation at 32°C after two days of ramping at 2°C per day. After 5 d (trial 1) or 7 d (trial 2), when the fragments at 32°C showed bleaching, images were captured (as in A, for trial 1) and colors quantified as described in Materials and Methods. Y-axis, arbitrary units (au) of final color intensities.

**Figure 5.**
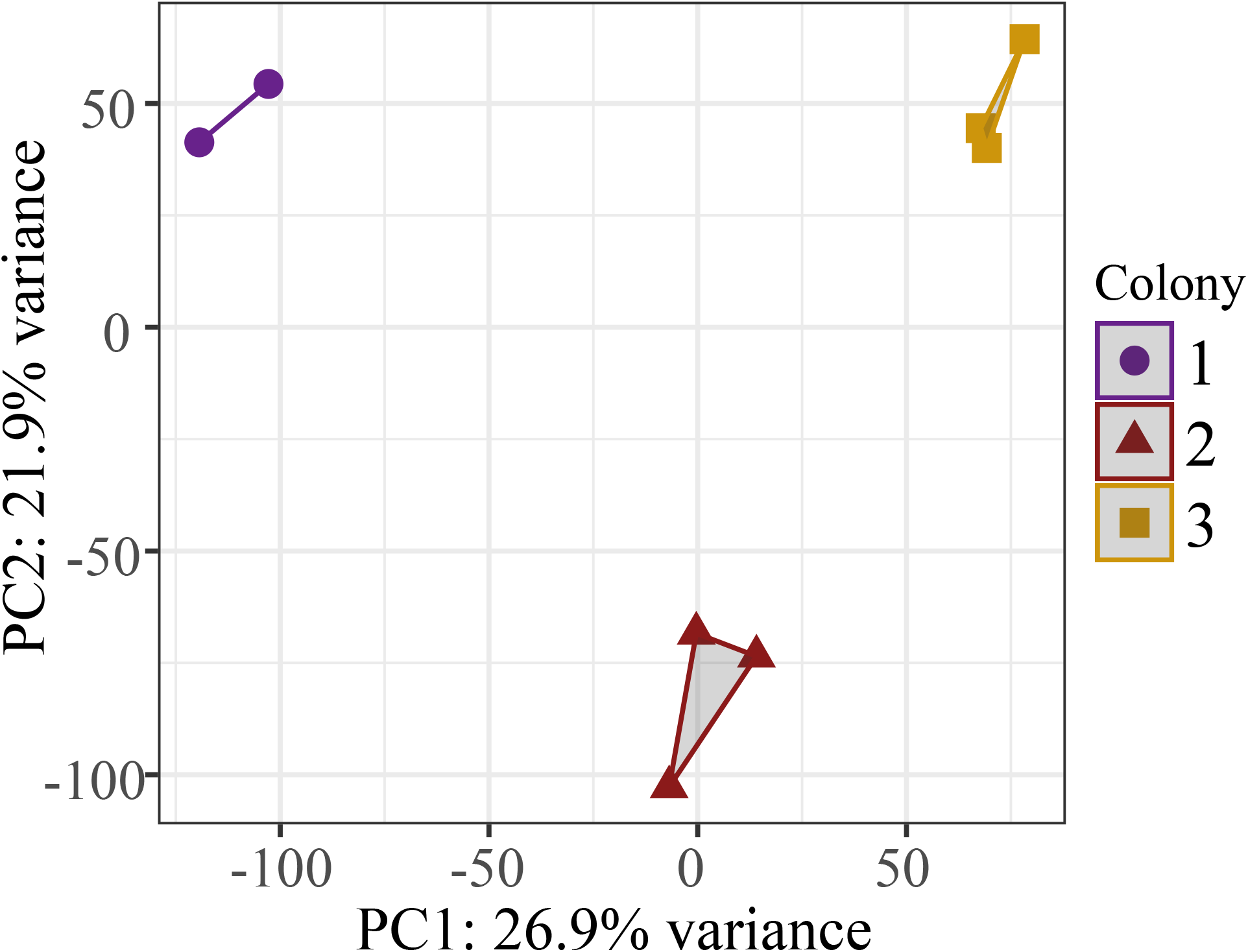
Unique gene expression profiles in *P. damicornis* colonies. Principal component analysis (PCA) of all r-log transformed isogroups clustered by *P. damicornis* colony (1, 2, or 3, as in Fig. 4A) demonstrating significantly different transcriptomic profiles during growth under control conditions.

NF-κB expression was quantified by Western blotting of lysates from the three *P. damicornis* colonies. Colony 3 (n=6) (hosting *Durusdinium* spp.) showed approximately 30-fold more NF-κB as compared to colonies 1 (n=5, P=0.015) and 2 (n=6, P<0.005; Figs. 4D, E) (hosting *Cladocopium* sp.).

We then asked if these distinct *P. damicornis* individuals showed differences in thermal tolerance by raising the temperature from 28°C to 32°C for ~1 week. Colony 3 showed much less bleaching than colonies 1 and 2 (Figs. 4A, F). Thus, the type of algal symbiont, the NF-κB levels, or both may influence the levels of resistance to bleaching under heat stress in this coral species.

### Genes possibly involved in thermotolerance in *P. damicornis*

The data above suggested a correlation between high levels of NF-κB and resistance of *P. damicornis* to heat-induced bleaching. To explore whether and how NF-κB might be involved in thermotolerance, we used RNA-seq to identify transcripts that were differentially expressed in colony 3 (thermotolerant, high NF-κB) vs. colonies 1 and 2 (thermosensitive, low NF-κB). We identified 466 genes whose expression was increased and 441 genes whose expression was decreased specifically in colony 3 (Files S6 and S7).

Because cnidarian NF-κB proteins can act as transcriptional activators (Wolenksi et al., 2011; Mansfield et al., 2017; Williams et al., 2018), we focused on the 116 annotated transcripts that were uniquely upregulated in colony 3 (Fig. 6). A GO enrichment analysis of these genes did not identify any over-represented biological processes, cellular components, or molecular functions. However, a literature-based search using the 116 annotated genes upregulated in colony 3 revealed that 15 (13%) have previously been reported to be upregulated during heat stress in some biological system, and 53 (46%) have been reported to be upregulated by some form of cellular stress (Table S2).

**Figure 6.**
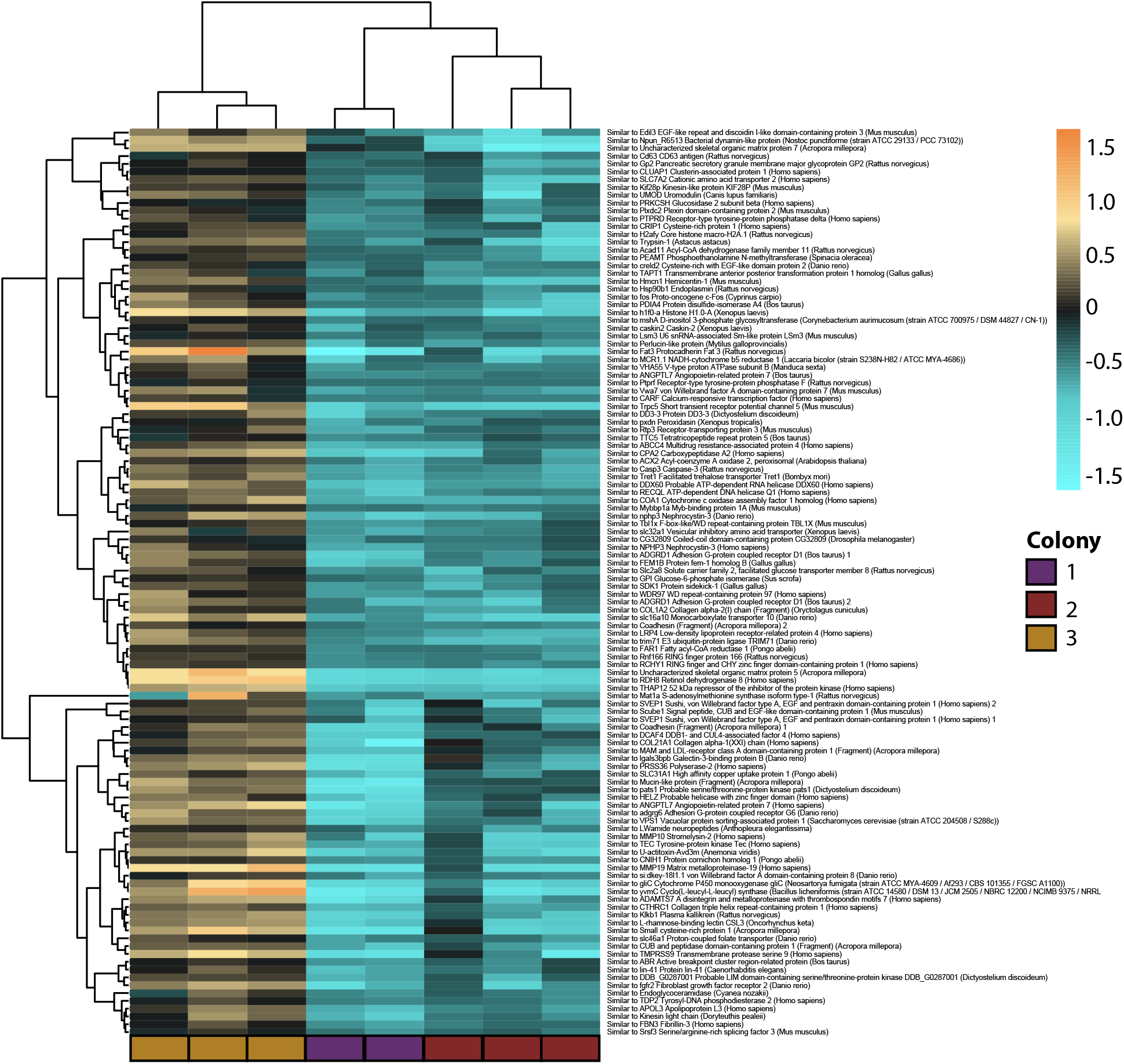
Genes upregulated in a thermoresistant *P. damicornis* colony 3. Heatmap for differentially expressed genes that were significantly upregulated in colony 3 relative to colonies 1 and 2. Only the 116 annotated genes (of 466 total – see File S6) were analyzed. Each row is a gene and each column is from an independent gene-expression library. The color scale is in log_2_ (fold change relative to a given gene’s mean expression), and genes and samples are clustered hierarchically based on Pearson’s correlation of their expression across samples. Block colors at the base of the heatmap indicate the colony of origin for each library. Hierarchical clustering of libraries (columns) showed statistically significant differences in gene expression across colonies.

Others have proposed that stress-response genes may be “frontloaded” in thermoresistant corals, thus providing protection upon the onset of stress (Barshis et al., 2013; Brener-Raffalli et al., 2018). Among the 116 annotated genes showing increased expression in colony 3, two (*creld2*, “cysteine-rich with EGF-like domains 2”, and von Willebrand factor A domain-containing 8) were reported previously to be frontloaded in wild populations of *Pocillopora* (Brener-Raffalli et al., 2018). Each of these genes has one predicted NF-κB binding site within 500 bp of its transcriptional start site (Fig. S3). Two additional genes (*Pd-cyst-rich* and *Pd-DPP7*) were previously reported to be substantially downregulated by heat-induced bleaching in *P. damicornis* (Vidal-Dupoil et al., 2009). Namely, in two-week bleaching experiments, *Pd-cyst-rich* was down-regulated by approximately 100-fold and *Pd-DPP7* was less robustly down-regulated (Vidal-Dupoil et al., 2009). Within 500 bp upstream of their transcriptional start sites, *Pd-cyst-rich* has two predicted NF-κB binding sites, and *Pd-DPP7* has three such sites (Fig. S3).

## Discussion

In this study, we have characterized the expression and activity of transcription factor NF-κB in two cnidarians for its possible role in symbiosis, immunity, and thermal tolerance. Although NF-κB has well-documented roles in immune- and stress-related processes in triploblasts, this is the first study to provide experimental evidence suggesting a role of NF-κB in resistance to bacterially induced disease and thermal stress-induced bleaching in cnidarians. Overall, our results suggest that the relationship between suppression of NF-κB during the establishment of symbiosis in Aiptasia larvae or the maintenance of symbiosis in adult Aiptasia is the result of a complex interaction between the symbiont and the host.

We previously showed that hosting of the algal symbiont SSB01 by Aiptasia larvae and adults results in reduced NF-κB expression as compared to the corresponding aposymbiotic anemones (Mansfield et al., 2017). In this report, we find that incubation of Aiptasia larvae with other Symbiodiniaceae genera (SSA01, SSA03, SSE01, Mf2.2b) does not reduce NF-κB expression (Figs. 1B, C). The failure of SSA03 and SSE01 to reduce NF-κB in larvae could be due to their inability to efficiently populate Aiptasia (Fig. 1A), but whether their inability to establish residency in Aiptasia larvae is due, at least in part, to their inability to downregulate NF-κB remains unclear. On the other hand, SSB01, SSA01, and Mf2.2b can all populate Aiptasia larvae (Hambleton et al., 2014; Wolfowicz et al., 2016; Fig. 1A), and yet only SSB01 downregulates NF-κB expression (Figs. 1B, C). Thus, it seems clear that suppression of NF-κB is not required by all Symbiodiniaceae genera in order to populate Aiptasia larvae.

How the presence of SSB01 in Aiptasia larvae leads to reduced NF-κB expression is not known. Our finding that incubation of aposymbiotic larvae with the human cytokine TGFβ1 also caused a reduction in NF-κB mRNA and protein staining (Fig. 1D-F) suggests that SSB01 reduces NF-κB by affecting a host factor(s) such as TGFβ. Consistent with that hypothesis, TGFβ has previously been shown to promote symbiosis in Aiptasia (Detournay et al., 2012) and in the coral *Fungia scutaria* (Berthelier at al., 2017). TGFβ comprises a superfamily of cytokines that induce anti-inflammatory and immune-suppressive effects in vertebrate systems (Travis and Sheppard, 2013; Chen and Ten Dijke, 2016), and the predicted Aiptasia TGFβ homologue is 51% identical to human TGFβ1 (Detournay et al., 2012). Taken together, these results support a model in which a host factor can suppress NF-κB which, in turn, facilitates the efficient establishment of symbiosis by certain algal symbionts, including SSB01 in Aiptasia.

Even though short-term infection of Aiptasia larvae with SSA01 does not reduce NF-κB expression, adult anemones that are stably populated by their endogenous algal symbionts (largely or entirely algae of strain SSA01) show low levels of NF-κB that are similar to the reduced levels seen in adults stably hosting SSB01. Furthermore, three *Durusdinium* isolates populate adult Aiptasia at a reduced level as compared to symbionts SSB01 and SSA01 (Fig. 2B; Matthews et al., 2018), and in no case did we find that *Durusdinium* sp. downregulated NF-κB. Thus, there are both native (SSA01) and non-native (*Durusdinium* spp.) Symbiodiniaceae that can establish symbioses with adult Aiptasia that do not appear to be able to reduce NF-κB expression or activity in short-term infections with Aiptasia larvae. It is notable that the three *Durusdinium* isolates that can form symbiosis with Aiptasia adults without reducing NF-κB levels (Fig. 2A) are maintained at lower levels than algal strains SSB01 and SSA01, which do reduce NF-κB levels in adults (Fig. 2B). It is not clear whether the inability of *Durusdinium* (a non-native symbiont) to reduce NF-κB causes it to be propagated at reduced levels in Aiptasia. Overall, our results show that different Symbiodiniaceae can have multiple and varying effects on the NF-κB signaling pathway, and suggest that different Symbiodiniaceae have different strategies for affecting host pathways in the establishment and maintenance of symbiosis.

Symbiotic Aiptasia hosting SSB01 have reduced NF-κB protein levels (Fig. 2A) and DNA-binding activity (Fig. 3A, B) as compared to the corresponding aposymbiotic Aiptasia. Moreover, after infection of Aiptasia with the bacterial pathogen *S. marcescens*, approximately 95% of aposymbiotic anemones (high NF-κB) survived whereas only 17% of symbiotic anemones (low NF-κB) survived (Fig. 3D). These results suggest a trade-off during symbiosis with adult Aiptasia hosting certain algal strains wherein NF-κB is reduced to facilitate the symbiotic partnership, but this reduction in NF-κB also results in increased susceptibility to pathogen-induced disease. Other host-microbe relationships that lead to reduced host NF-κB activity include Drosophila-gut microbiota (Chu and Mazmanian, 2013), salamander-algae (Burns et al., 2017), honeybee-deformed wing virus (Nazzi et al., 2012; Di Prisco et al., 2016), and bobtail squid-*V. fischeri* (Chun et al., 2008).

We have also found that three colonies of the coral *P. damicornis* that have been maintained in common garden conditions for >5 years can 1) have differential bleaching susceptibilities in response to increased water temperature, 2) harbor different Symbiodiniaceae genera, and 3) have different basal levels of NF-κB (Figs. 4A-F). The most thermoresistant colony (#3) had constitutively higher NF-κB levels and hosted a variety of *Durusdinium* symbionts as compared to thermosensitive colonies 1 and 2, which had lower NF-κB levels and hosted primarily *Cladocopium* symbionts. Several studies have reported that hosting *Durusdinium* is associated with increased thermotolerance in corals (e.g., Barshis et al., 2013; Silverstein et al., 2017; Stat and Gates, 2011, Davies et al., 2018). Whether host factors, such as increased NF-κB levels, also contribute to the bleaching resistance in our *P. damicironis* colony 3 is not clear at this time. However, it is notable that the stable hosting of *Durusdinium* species does not reduce NF-κB levels during symbiosis in both Aiptasia (Fig. 2A) or *P. damicornis* (Fig. 4D, E). Therefore, cnidarians that naturally host *Durusdinium* may have increased thermoresistance to bleaching due to the following: 1) an intrinsic thermoresistance of *Durusdinium* symbionts; 2) effects of *Durusdinium* on host physiology (e.g., by maintaining high NF-κB levels); or 3) a combination of both. Relevant to these three possibilities, *Durusdinium* does not appear to be inherently more thermoresistant than other Symbiodiniaceae genera when grown *in vitro* in the absence of a cnidarian host (Chakravarti and van Oppen, 2018), suggesting that effects of *Durusdinium* on host physiology are important for thermotolerance.

We identified 116 annotated genes with significantly higher expression levels in thermoresistant *P. damicornis* colony 3 as compared to thermosensitive colonies 1 and 2. Several groups (e.g., Barshis et al., 2013; Brener-Raffalli et al., 2018) have proposed that higher basal expression of certain host stress response genes, i.e. “frontloaded genes”, is associated with protection against thermally induced bleaching in reef-building corals. Thus, the upregulation of certain genes in our *P. damicornis* colony 3 could be indicative of frontloaded stress-response genes. Consistent with that, we found that approximately 46% of the 116 colony 3-specific upregulated genes have been reported to be upregulated by stress in other organisms (Table S2). Moreover, two of the colony 3-specific upregulated genes (*creld2* [cysteine-rich with EGF-like domain 2-like] and von Willebrand factor A domain-containing 8) were also reported as frontloaded genes in wild populations of bleached *Pocillopora* (Brener-Raffalli et al., 2018). Of note, the *creld2* and von Willebrand factor A domain-containing 8 genes both have predicted NF-κB binding sites near their transcriptional start sites (Fig. S3).

It is likely that many of the genes that show differences in basal expression in colony 3 vs. colonies 1 and 2 are not a consequence of increased NF-κB expression or increased thermoresistance in colony 3. For example, Granados-Cifuentes et al. (2013) have shown that there are extensive differences in gene expression among genetically distinct *Acropora millepora* nubbins, even when these coral are raised in common garden lab conditions for four weeks. In addition, colony 3 harbors a different genera of symbiont (i.e., *Durusdinium*) than colonies 1 and 2 (i.e., *Cladocopium*), and algal symbiont type has been shown to affect host gene expression patterns (DeSalvo et al., 2010; Barshis et al., 2013). Others have proposed that higher plasticity in gene expression (rather than frontloading or symbiont type) can also enable coral to more effectively resist thermal stress (Granados-Cifuentes et al., 2013; Seneca and Palumbi, 2015; Davies et al., 2016; Kenkel and Matz, 2016).

The frontloading and plasticity gene expression models of thermoresistance in corals both consider only genes that respond to or protect against the stress of increased temperature. However, others (Dixon et al., 2015) have seen a negative correlation of certain heat tolerance-associated genes with bleaching susceptibility. Thus, we propose that there is another class of genes that are involved in bleaching susceptibility: namely, genes whose upregulation or downregulation *is required for* thermal bleaching. In this scenario, one might predict that bleaching effector genes that are normally upregulated during thermal stress would be basally lower in thermoresistant corals, and that bleaching effector genes that are downregulated during thermal stress would be basally higher in thermoresistant corals. Thus, in thermoresistant colony 3, NF-κB could be transcriptionally activating some genes that must be sufficiently downregulated by heat in order for bleaching to ensue. Related to this hypothesis, we find that the basal levels of two genes (*Pd-cyst-rich* and *Pd-DPP7*), which have been reported as highly downregulated in bleached *P. damicornis* (Vidal-Dupoil et al., 2009), are overexpressed in thermoresistant colony 3 as compared to thermosensitive colonies 1 and 2, and both genes have multiple predicted NF-κB binding sites within 500 bp of their transcriptional start sites. Although there has been an amazing lack of consistency among gene expression studies that have compared mRNA levels in control vs. bleached corals, genes encoding cysteine-rich proteins have been identified as being downregulated in at least two other studies that have looked at genes affected by bleaching in the corals *P. damicornis* (Vidal-Dupoil et al., 2009) and *Orbicella faveolata* (DeSalvo et al., 2008).

Many studies that have looked at gene expression changes that occur during bleaching or that are associated with thermoresistance have used GO (Gene Ontology) analysis in an attempt to identify biological processes that are affected by bleaching or heat. Nevertheless, such studies provide correlations with mRNA expression levels, and not with protein activity. Furthermore, such studies are inherently compromised by a lack of proven gene homologues among the datasets analyzed, as has been suggested by others (Djordevic et al., 2016). Therefore, we propose that studies, such as ours and a recent study of the c-Jun N-terminal kinase JNK (Courtial et al., 2017), that are investigating induced or inherent differences in transcription factor activity will ultimately be required to identify cell networks and pathways that contribute to biological processes related to symbiosis and thermal sensitivity.

## Data accessibility

Raw Fastq files are available via Sequence Read Archive via accession codes PRJNA514346 for ITS2 and PRJNA514335 for 18S. Raw fastq files for gene expression work can be accessed via Sequence Read Archive code PRJNA521995. File sets 1-9 are available upon request at

## Authors’ Contributions

T.D.G. supervised the overall project. K.M.M, P.A.C., S.W.D., and T.D.G. conceived and designed the experiments. K.M.M performed experiments except the following; P.A.C. (Aiptasia spawning, larval symbiont infections, and adult Aiptasia Western blotting and quantification); P.A.C and D.J.C (generation and analyses of adult Aiptasia harboring Mf2.2b, Ap02, A001 algal strains); E.V.V. and M.P. (*S. marcescens* infections of Aiptasia); N.G.K. (*P. damicornis* sample and data collection, Symbiodiniaceae ITS2 and coral 18S sequencing) B.E.B. (*P. damicornis* maintenance, experimental setup, sample and data collection, and tagseq library preparation); O.H., (*P. damicornis* maintenance, sample and data collection, colorimetric assay); T.W.S (computational analysis of promoter sequences in the Aiptasia genome); J.R.P (supervision of larvae experiments); S.W.D. (supervision and experimental design of *P. damicornis* experiments as well as tagseq data analysis). K.M.M, P.A.C., N.G.K., B.E.B., J.A.P., S.W.D., and T.D.G wrote the paper.

## Acknowledgements

We thank Kim Ritchie (University of South Carolina Beaufort) for *S. marcescens*, Virginia Weis (Oregon State University) for several Aiptasia lab strains, Mary Alice Coffroth for Mf2.2b, A001, and Ap2 algal strains, Justin Scace for the *P. damicornis* samples, Rachel Wright for help with the *P. damicornis* experiment, Nikki Traylor-Knowles for providing access to the *P. damicornis* genome, Todd Blute for help with microscopy, and Ulla Hansen, Albert Mondragon, Sean Mullen, and Leah Williams for helpful discussions.

