## Supplemental Materials for "Varied effects of algal symbionts on transcription factor NF-κB in a sea anemone and a coral: possible roles in symbiosis and thermotolerance"

**Supplementary Files**

File S1. Raw count data for differential expression analysis
File S2. ITS2 count data for *P. damicornis*
File S3. Genes significantly upregulated in colony 1 relative to colonies 2 and 3
File S4. Genes significantly downregulated in colony 1 relative to colonies 2 and 3
File S5. Genes significantly upregulated in colony 2 relative to colonies 1 and 3
File S6. Genes significantly downregulated in colony 2 relative to colonies 1 and 3
File S7. Genes significantly upregulated in colony 3 relative to 1 and 2
File S8. Genes significantly downregulated in colony 3 relative to 1 and 2
File S9. List of NF-κB binding sites from Aiptasia PBM assay used for promoter analysis

#### Supplemental Tables

**Table S1. Primers for qPCR used on Aiptasia.**

| | |
|---|---|
| Aiptasia NF-κB gene | 5'- ATTTTGGCAGTGGTTTCCTG<br>5'- TGGCAGCATAGTCTTGCATC |
| Aiptasia RPL10 gene (from Poole et al., 2016) | 5'- ACGTTTCTGCCGTGGTGTCCC<br>5'- CGGGCAGCTTCAAGGGCTTCA |

**Table S2. Genes significantly upregulated in colony 3 that have been associated with stress and physiological responses.**

| *Gene* | *Organism* | *Inducer* | *Reference* | *PubMed link* |
|---|---|---|---|---|
| CLUAP1 | Human | Heat | Wang et al, 2016 | www.ncbi.nlm.nih.gov/pubmed/26853533 |
| SLC7A2 | Pig | Heat | Cervantes et al, 2016 | www.ncbi.nlm.nih.gov/pubmed/27264891 |
| PRKCSH | Plant | Heat | Nwugo et al, 2016 | www.ncbi.nlm.nih.gov/pubmed/27842496 |
| HSP90 | Many | Heat | Chen et al, 2018 | www.ncbi.nlm.nih.gov/pubmed/29920826 |
| c-Fos | Human | Heat | Andrews et al, 1987 | www.ncbi.nlm.nih.gov/pubmed/3601676 |
| PDIA4 | Fish | Heat | Mahanty et al, 2016 | www.ncbi.nlm.nih.gov/pubmed/27058960 |
| Perlucin-like | Mussel | Heat | Fields et al, 2016 | www.ncbi.nlm.nih.gov/pubmed/27335449 |
| TRPC5 | Human | Heat | Baez et al, 2014 | www.ncbi.nlm.nih.gov/pubmed/25366233 |
| TTC5 | Human | Heat | Davies et al, 2015 | www.ncbi.nlm.nih.gov/pubmed/21147850 |
| Casp3 | Pufferfish | Heat | Cheng et al, 2018 | www.ncbi.nlm.nih.gov/pubmed/29276954 |
| Fem1B | Red flour beetle | Heat | Xiong et al, 2018 | www.ncbi.nlm.nih.gov/pubmed/28681272 |

| | | | | |
|---|---|---|---|---|
| gliC | Whitefly | Heat | Guo et al, 2018 | www.ncbi.nlm.nih.gov/pubmed/30195394 |
| Klkb1 | Human | Heat | Joseph et al, 2002 | www.ncbi.nlm.nih.gov/pubmed/12489799 |
| CSL3 | Coral | Heat | Zhou et al, 2017 | www.ncbi.nlm.nih.gov/pubmed/28069433 |
| FGFR2 | Pig | Heat | Rogers et al, 2008 | www.ncbi.nlm.nih.gov/pubmed/18988085 |
| CD63 | Human | Oxidative stress | Sheller et al, 2016 | www.ncbi.nlm.nih.gov/pubmed/27333275 |
| PXDN | Human | Oxidative stress | Hanmer & Mavri-Damelin, 2018 | www.ncbi.nlm.nih.gov/pubmed/29953917 |
| Scube | Rat | Oxidative stress, ischemia | Turkmen et al, 2013 | www.ncbi.nlm.nih.gov/pubmed/23517257 |
| VPS1 | Yeast | Oxidative stress | Mikawa et al, 2010 | www.ncbi.nlm.nih.gov/pubmed/20070859 |
| ANGPTL7 | Human | Hypoxia | Parri et al, 2014 | www.ncbi.nlm.nih.gov/pubmed/24903490 |
| GPI | Human | Hypoxia | Minchenko et al, 2017 | www.ncbi.nlm.nih.gov/pubmed/29236388 |
| SRSF3 | Human | Hypoxia | Brady et al, 2017 | www.ncbi.nlm.nih.gov/pubmed/28961236 |
| KIF28P | Yeast | Salt stress | Schoch et al, 1997 | www.ncbi.nlm.nih.gov/pubmed/9371882 |
| UMOD | Mice | Salt stress | Graham et al, 2018 | www.ncbi.nlm.nih.gov/pubmed/30216136 |
| PEAMT | Plant | Salt stress | Tabuchi et al, 2005 | www.ncbi.nlm.nih.gov/pubmed/15695433 |
| CRELD2 | Mice | ER Stress | Oh-Hashi et al, 2018 | www.ncbi.nlm.nih.gov/pubmed/30111858 |
| COA1 | Human | Metabolic stress | Zhang et al, 2016 | www.ncbi.nlm.nih.gov/pubmed/27550821 |

| | | | | |
|---|---|---|---|---|
| SDK1 | Human | Serum starvation | Yoon et al, 2012 | www.ncbi.nlm.nih.gov/pubmed/2210536 |
| TRET1 | Human | Freezing | Uchida et al, 2017 | www.ncbi.nlm.nih.gov/pubmed/28552273 |
| PTPRF | Mice | Diesel exhaust | Goodson et al, 2017 | www.ncbi.nlm.nih.gov/pubmed/28751527 |
| TBL1X | Human | Smoke | Katbamna et al, 2013 | www.ncbi.nlm.nih.gov/pubmed/23665419 |
| Hmcn1 | Oyster | Zinc | Luo et al, 2017 | www.ncbi.nlm.nih.gov/pubmed/28304108 |
| TDP2 | Plant | Copper | Macovei et al, 2010 | www.ncbi.nlm.nih.gov/pubmed/20458495 |
| Far1 | Wheat | Fosthiazate | Yin et al, 2012 | www.ncbi.nlm.nih.gov/pubmed/23520855 |
| RECQL | Human | DNA damage | Popuri et al, 2012 | www.ncbi.nlm.nih.gov/pubmed/23095637 |
| PLXDC2 | Human | Pacitaxel | Wang & Li, 2018 | www.ncbi.nlm.nih.gov/pubmed/29928353 |
| CRIP1 | Human | UV, staurosporine | Latonen et al, 2008 | www.ncbi.nlm.nih.gov/pubmed/18177859 |
| ABCC4 | Human | Cisplatin | Savaraj et al, 2003 | www.ncbi.nlm.nih.gov/pubmed/12792791 |
| RCHY1 | Human | Arsenic trioxide | Yan et al, 2014 | www.ncbi.nlm.nih.gov/pubmed/25116336 |
| COL21A1 | Human | Chemotherapeutic drugs | Januchoswki et al, 2016 | www.ncbi.nlm.nih.gov/pubmed/27390605 |
| Caskin2 | Fish | 17α-methyltestosterone | Gao et al, 2017 | www.ncbi.nlm.nih.gov/pubmed/?term=Caskin-2 |
| COL1A2 | Human | Perfenidone | Hisatomi et al, 2012 | www.ncbi.nlm.nih.gov/pubmed/22694981 |
| slc16a10 | Rabbit | Low thyroid hormone | Mebis et al, 2009 | www.ncbi.nlm.nih.gov/pubmed/19439506 |

| | | | | |
|---|---|---|---|---|
| Mat1a | Human | S-adenosyl-L-methionine | Lozano-Sepulveda et al, 2016 | www.ncbi.nlm.nih.gov/pubmed/27076759 |
| SVEP1 | Human | TNF; estradiol | Glait-Santar & Benayahu, 2012 | www.ncbi.nlm.nih.gov/pubmed/22265959 |
| lgals3bpb | Human | TNF; EBV/lupus | Rasmussen et al, 2016; Noma et al, 2012 | www.ncbi.nlm.nih.gov/pubmed/27084029 www.ncbi.nlm.nih.gov/pubmed/22447108 |
| CUB serine protease | Shrimp | LPS,PGN,β-glucan | Yang et al, 2017 | www.ncbi.nlm.nih.gov/pubmed/28619282 |
| MMP19 | Human | Wound healing; Anisomycin | Mendoza-Garcia et al, 2015; Sampieri et al, 2008 | www.ncbi.nlm.nih.gov/pubmed/26094764 www.ncbi.nlm.nih.gov/pubmed/18029162 |
| ADAMTS7 | Human | Aortic aneurysm | Qin et al, 2017 | www.ncbi.nlm.nih.gov/pubmed/28849199 |
| CTHRC1 | Human | Aortic valve disease | Liu et al, 2017 | www.ncbi.nlm.nih.gov/pubmed/29212967 |
| APOL3 | Human | HCV infection | Shrivastava et al, 2016 | www.ncbi.nlm.nih.gov/pubmed/27193023 |

**Supplemental Figures**

**Figure S1**

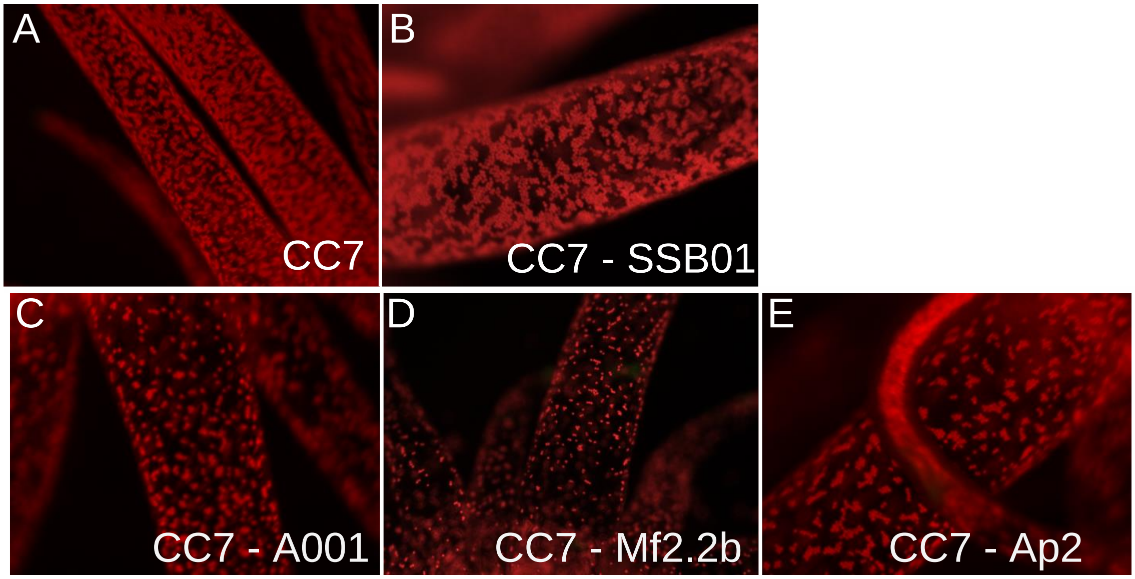


**Figure S1. Aiptasia symbiotic with different isolates of Symbiodiniaceae.** Representative micrographs of algal chlorophyll autofluorescence within adult CC7 anemone tentacles symbiotic with the following: (A) their endogenous algae, (B) SSB01, (C) A001, (D) Mf2.2b, and (E) Ap2. The adult anemones have been in stable symbiosis under standard growth conditions for > 1 year.

**Figure S2**

# A

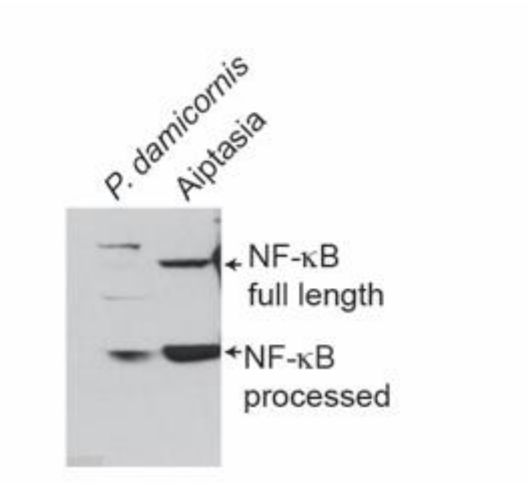


# B

Range 1: 3 to 750 Graphics Next Match Previous Match

| Score | Expect | Method | Identities | Positives | Gaps |
|---|---|---|---|---|---|
| 881 bits(2277) | 0.0 | Compositional matrix adjust. | 445/757(59%) | 553/757(73%) | 28/757(3%) |

```
Query  2      THSEQQVGGTLTDSMLHEIMTPGFLPDISSLNVQMEMGYEGPYLEILEQPKSRGFRFRYP  61
              T+SE+Q+G TLTDS + +++TPG+LPDIS+L V     Y+GPY+EILEQPK RGFRFRYP
Sbjct  3      TNSERQIGETLTDSFVMDLLTPGYLPDISALQVPTA-SYQGPYMEILEQPKQRGFRFRYP  61

Query  62     CEGPSHGGLPGEFSDSKNKSYPSVQVCNYQGPCRIVVSLVTEDEPHMPHAHSLTGKHANN  121
              CEGPSHGGLPG++S+   KSYPSVQ+CNYQGP RIVVSLVT DEP MPHAHSL  K++NN
Sbjct  62     CEGPSHGGLPGQYSEKGKKSYPSVQLCNYQGPARIVVSLVTVDEPPMPHAHSLISKNSNN  121

Query  122    DGIVTVQIGTEQGMTASFPNLGIQHVTKKKVAKTLTERYTKMQALQNATLTALATNNSTP  181
               G+VTVQIG EQGMTA+FPNLGI+HVTKK V+K L +RY KMQ L  ATL AL + +
Sbjct  122    -GVVTVQIGPEQGMTATFPNLGIEHVTKKMVSKVLMDRYIKMQTLHTATLNALTSGDG--  178

Query  182    SSFMNFGSVAREQVMASQGPFDRNLAAAVAGEETKKILKLVQEQSKTMNLSAVRLCFQAY  241
                F   G V +  V   +G FD+ LA AVA EE++K+  +V+EQ ++MNL+AVRLCFQA+
Sbjct  179    KVFGVAGLVDQAMVDGDRGSFDKRLAEAVAEEESQKVRAMVEEQKQSMNLNAVRLCFQAF  238

Query  242    LPDENGNFTKPLKPCISNPVYDSKAPASCQLKICRMDKNSGCVTGGDEIYLLCDRVQKDD  301
              LPDE G FTK L PCISN VYDSKAP++  LKICRMD+NSGCV GGDE+YLLCD+VQKDD
Sbjct  239    LPDETGAFTKALPPCISNAVYDSKAPSASNLKICRMDRNSGCVKGGDEVYLLCDKVQKDD  298

Query  302    IEIRFYENN-DDGKPIWEDTGKFAPADVHRQFAIVFKTPAYHNIAIERPVEVLLELRRKS  360
              IE+ FYE   D GK  WED G F+P DVHRQ AIVFKTP Y N+AIE+PV+V LELRRKS
Sbjct  299    IEVIFYETEMDTGKKTWEDRGVFSPTDVHRQVAIVFKTPPYWNVAIEQPVKVQLELRRKS  358

Query  361    DKETSEPFTFTYSPQMFDTEQIGAKRQKKVPHFSDYYPPGGPPGAAGGGGGGFNFGS---  417
              D+ETS+P  FTY PQMFD EQIGAKR+KK+PHFSDY   GG  G   G GG    G
Sbjct  359    DQETSDPVEFTYQPQMFDNEQIGAKRRKKIPHFSDYLGGGGGGGGGPGMGGAGGGGGGFN  418
```

**Figure S2. Antiserum against Aiptasia NF-κB cross-reacts with NF-κB from *P. damicornis*.** (A) Extracts from symbiotic *P. damicornis* and Aiptasia were analyzed by anti-NF-κB Western blotting using our anti-Aiptasia NF-κB antiserum that was generated against the Rel Homology Domain (RHD) of NF-κB (Mansfield et al., 2017). (B) Protein Sequence alignment of the Rel Homology Domain (RHD) of Aiptasia NF-κB (Query) and *P. damico*rnis NF-κB (Sbjct).

Alignment Generated using NCBI's protein blast align function (https://blast.ncbi.nlm.nih.gov/Blast.cgi).

### Figure S3

## A

**Sequences 500 bp upstream of the start sites of transcription for cysteine-rich with EGF-like domain 2-like and von Willebrand factor A domain-containing 8 genes.**

isogroup 00012457 *creld2 Cysteine-rich with EGF-like domain 2-like* (Brener-Raffalli, bioRxiv posted 2018)
1 site:
CGGTTTTCCC -51 to -42 (z-score 3.989154339)

-500
AGATGTAAATGACAGCAGTACCGCAAAATCTGTGCTAAACTTTGAAGTGCAAGGAAGTACTATGCAGCT
TTCAATCGATCCAAACCAACCAAAATTTTCCACGGCCAAAGCTCGTGTCTTGTTCTAATTGGCTAATCTTA
GTCACGTGCTTAACGTGGAGCCCTTTGGTTGGTTTATTTTTCCATGTCATTTCATTGATTCCCATGAGTGA
GCCCTCGCTTTGCGCCGAGGTTTAAAAAGTATTTGAGAGAATTTACTTTTAAACTTATTAAAAAGCTACTT
AGCTTTGTTCTCAAATGATCAATAGATCTACCAACTCTGAAAATTTCAGATATATTTTAATGTTTTGTCTCA
CCTCACGTGGAATGATTATGAATTTAGGACGCAAAAAAGTGATACAAATTATTGTACAATGGCTGATCA
AAACTTTATTCTCCTATCTATTAGCGCGGTTTTCCCTGACTCACCAGCGGTTTTCTCTTCAAGAGAAACAC
GATAAT -1

isogroup00020032 *von Willebrand factor A domain-containing 8* (Brener-Raffalli, bioRxiv posted 2018)
1 site:
TAAAAATCCC -132 to -123 (z-score 5.655)

-500
TATTTTACTCTTAGTGTGAGTATCATTTCCCGAAAAACGTTGGCGAAGGAACTGCTCTTTCGTAGTTGTTA
GCAGGGACTTCAAATTTTAAATTTGTTGCACCTACAATGGATCTATGTTTCAGCGAAAATAATCATACATT
CAAGAGTAAACAATGTAGAAAAGGCCAATATTTCAATGGAGTATAAAGTCACGCTAAATTCACCAGTGA
CGCGTGTCAAAATATTGAACGAGTTAGGAGGCAGAGTGGGCTCCTGACAAGTAATGTAATTTCTCTCGG
TATACCATAAAGCCTTTAAATAAATATAAAAACAAACAAAGCAAAACAAAACAAAATAATCTAGTGTACG
AATAGACATAAAACAATATAAAAATCCCAACTACCACCCTACGTCAACACATCGAGGAATGCGATTTAGA
AAGTGTAAACGAGCAAAGGAGAAATAGAATAATGAGAAAAGTAGGTTCCAGTCTTTCACTGATTTCAAG
CGATCCATCAT -1

## B

**Sequences 500 bp upstream of the start sites of transcription for *Pd-cyst-rich* and *Pd-DPP7***

Isogroup00006164 *Pd-cyst-rich* (Vidal-Dupiol et al., 2009)
2 sites:
GGTCTTCCCC -373 to -364 (z-score 5.91)
AATTCCCTTT -275 to -266 (z-score 3.20)

-500
CTTAATAAATTTGCATTTTTGAGCAAGTTTTTTGAAACAAACTTTATAATCATTTAATGATCAATTATGGCA
TAAAATAGCGATTTTTCGTCGATTTAAACCCCATTCTGTGGAATTTGTATTGAGTGGTCTTCCCCAAGGGT
GACCACTTAATACAGGTTTGATTGCAGTTCTTGGCCATATTTTGTTTTTCGGTCTCGTTATTATCTTCTAAT
CATAAACTGCAATTCCCTTTTAATCTTTCTCATCCTAGGCCGTATTTTTCCATGTCAATTCAGTAATTTCCG
ATTGACTCTTCTTTTTTCCTGTTCTAAAGGAAACGATGCTGATGTTGCAATACATTATGCTGAATAAATTTT
CCTCGTCACGTAAAAACAACAAAAAAAAATCATTTAACCTGTAAAGGTCAGTGATGACGGCATCGACAG
CACGCTTTAGTTTCAATCCCTCAGTAATGATCGTGACGACAACAATTTCAGTCACGTGGTCTGTCCAATG
AG -1

isogroup00024800 *Pd-DPP7* (Vidal-Dupoil et al., 2009)
3 sites:
GGGGGAGGAA -484 to -475 (z-score 5.20)
GGGGACACCC -414 to -405 (z-score 5.89)
CCCCTCCCCC -286 to -276 (z-score 4.84)

-500
AGGGGGTGGACTGGGTGGGGGAGGAAAGCCTTGGAAAGGATCACAGAAAATCCTATATGGCTGCATG
CCTATCTTAGAGGGGAAGAGGGGACACCCATAGGAAACAATTTCAATCTGCAACTGATTGCCTATGTAC
TATAGACTTGTGAAGTAAAAAATAGCTTTGAATTGAAGCCTGGAATCGAAGTAAGTCAAGAACCAGGTG
CTGAATGCACCCCTCCCCCCCTCCCCCCCAACAAACAGCATTTGCAGCAGTATCTTTAAAGGTTGGTTAC
TGGAGGTCTATCGCTGCCCATAAGACCTCTTACGCCTGTTACACAGCTGTCTCTTACACCATCAGCAGCC
ACATATGGAATAAGTTATTGAAACCCCATGCCAGTATTTAGAATTTGTCTTTCCTGAGGCCTTGAGACAC
CAGTTTTGGGGCCTGTTTCCTGAGGATGGCTTGTGATACTTGGCAAGAACATGGAGGTGTTCTGAGTGG
AATAATTTGTTAAGC -1

**Figure S3. Predicted NF-κB binding sites in promoter sequences of genes possibly contributing to thermal resistance or bleaching in *P. damicornis* colony 3.** Shown are sequences located 500 bp upstream of the transcriptional start sites of the indicated genes. Sequences are presented from positions -500 to -1, relevant to the transcriptional start site. Sequences were obtained from the *P. damicornis* genome (Cumming et al., 2018). Sequences in red are predicted NF-κB binding sites based on Protein Binding Microarray (PBM) analysis of Aiptasia NF-κB (see Mansfield et al., 2017, and File S9). The locations of the predicted NF-κB binding sites (relative to position -1) and their z-scores from the PBM analysis are indicated. (A) Possible frontloaded genes. (B) Possible bleaching effector genes. See text for more details on these genes.
